# DEsingle for detecting three types of differential expression in single-cell RNA-seq data

**DOI:** 10.1101/173997

**Authors:** Zhun Miao, Ke Deng, Xiaowo Wang, Xuegong Zhang

## Abstract

**Summary:** The excessive amount of zeros in single-cell RNA-seq data include “real” zeros due to the on-off nature of gene transcription in single cells and “dropout” zeros due to technical reasons. Existing differential expression (DE) analysis methods cannot distinguish these two types of zeros. We developed an R package DEsingle which employed Zero-Inflated Negative Binomial model to estimate the proportion of real and dropout zeros and to define and detect 3 types of DE genes in single-cell RNA-seq data with higher accuracy.

**Availability and Implementation:** The R package DEsingle is freely available at https://github.com/miaozhun/DEsingle and is under Bioconductor’s consideration now.

**Supplementary information:** Supplementary data are available at *bioRxiv* online.

## 1 Introduction

Single-cell RNA-seq (scRNA-seq) data have different characteristics from bulk RNA-seq data (Brennecke, et al., 2013). One important difference is that there are much more zero values in scRNA-seq data (Bacher and Kendziorski, 2016; Miao and Zhang, 2016). There are both biological and technical reasons. On the technical aspect, usually the amount of mRNAs in one cell is very tiny (∼0.01-2.5pg) (Boon, et al., 2011), and most genes have only small mRNA copy numbers in a cell (Macaulay and Voet, 2014; Marinov, et al., 2014). Due to the low efficiency of the mRNA capture protocol (Islam, et al., 2014; Marinov, et al., 2014), the mRNAs of some genes are totally missed during the reverse transcription and cDNA amplification steps, and consequently missed in the sequencing (Wang and Navin, 2015). This phenomenon is called “dropout” events (Kharchenko, et al., 2014). We call this type of zero values in the data as dropout zeros as they do not reflect the true expression status. On the other hand, because of the stochastic nature of transcription in single cells and the heterogeneity between cells, there is also a high chance that the expression levels of some genes are really zero in some cells at the time when they are sequenced (Delmans and Hemberg, 2016). We call this type of zero values as “real” zeros because they reflect the true status of the genes’ transcription. The excessive zero values observed in scRNA-seq data are the mixture of these two possible types of zeros. Discriminating these two types of zeros is important for downstream analyses. Existing methods for analysing differential gene expression in single cells did not take the different types of zeros into consideration.

We developed an R package DEsingle to define and detect 3 types of DE genes between two groups of scRNA-seq data. It employed Zero-Inflated Negative Binomial model to estimate the proportion of real and dropout zeros. Simulation experiments showed that DEsingle can detect DE genes with better overall performance than existing methods. We applied DEsingle on a scRNA-seq dataset of human preimplantation embryonic cells (Petropoulos, et al., 2016) and found that the 3 types of DE genes imply different biological functions. DEsingle is not only a new tool for detecting DE genes with high accuracy, but also provides a new view of DE genes by discriminating 3 types of differential expression.

## 2 Materials and methods

### 2.1 The ZINB model for discriminating the proportion of real and dropout zeros

DEsingle adopted the Zero-Inflated Negative Binomial (ZINB) model to describe the read counts and the excessive zeros in scRNA-seq data (Supplementary Fig. S1). The ZINB distribution is a mixture of constant zeros and a Negative Binomial (NB) distribution with a mixing parameter *θ* (Garay, et al., 2011). The probability mass function (PMF) of ZINB distribution for read counts *N*_*g*_ of gene *g* in a group of cells is

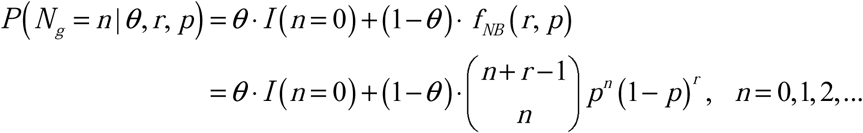

where *θ* is the proportion of constant zeros of gene *g* in the group of cells, *I*(*n* = 0) is an indicator function which equals to 1 for *n* = 0 and 0 for *n* ≠ 0, *f* _*NB*_ is the PMF of the NB distribution, *r* is the size parameter and *p* is the probability parameter of the NB distribution. The NB part can also have zero values. The observed zero values are the mixture of constant zeros and zeros from the NB distribution. We proved that this model can separate observed zeros as two parts to reflect the proportion of real and dropout zeros (see Supplementary Materials). Based on this, we can distinguish 3 types of differential expression according to whether the difference is in the zero part or the NB part of the distribution, or in both.

### 2.2 The Workflow of DEsingle

DEsingle uses the ZINB model to detect differentially expressed genes. It includes 3 major steps: data normalization, detection of DE genes, and subdivision of DE genes into 3 types (Supplementary Fig. S1E). The input of DEsingle is the raw read-count matrix from scRNA-seq data. DEsingle integrates a modified median normalization method similar to the one used in DESeq (Anders and Huber, 2010). After normalization, maximum likelihood estimation (MLE) and constrained MLE of the parameters of two ZINB populations are calculated. Finally, DEsingle uses likelihood-ratio tests to compare two ZINB populations to detect DE genes and to subdivide them into the 3 types: DEs, DEa and DEg. DEs refers to “different expression status”. It is the type of genes that show significant difference in the proportion of real zeros in the two groups, but do not have significant difference in the other cells. DEa is for “differential expression abundance”, which refers to genes that are significantly differentially expressed between the groups without significant difference in the proportion of real zeros. DEg or “general differential expression” refers to genes that have significant difference in both the proportions of real zeros and the expression abundances between the two groups. The detailed definition of the 3 types and description of the package are given in the Supplementary Materials.

## 3 Results and discussion

### 3.1 Performance evaluation on simulated data

We compared DEsingle with 7 existing methods on 8 simulated datasets that considers both stochasticity and mRNA capture efficiency (see Supplementary Materials). Supplementary Fig. S2 shows the ROC curves and the AUCs of the different methods. DEsingle performed the best on 7 of the datasets and was very close to the best on the other dataset.

### 3.2 Three types of DE genes found in real data experiments

We applied DEsingle on a human preimplantation embryonic cell scRNA-seq dataset (Petropoulos, et al., 2016). Results show that DEsingle can detect genes and pathways that have significant changes in their expression between days of embryonic cell development, and the 3 types of detected DE genes are enriched in different biological processes (Supplementary Fig. S3, S4). Distinguishing DE genes as 3 subtypes provides the extra possibility of analysing the expression patterns and functions in different cell groups. Details of the results and functional analyses are given in the Supplementary Materials.

## 4 Conclusion

We developed an R package DEsingle for differential expression analysis of scRNA-seq data. By estimating the proportion of real and dropout zeros with ZINB model, DEsingle not only detects DE genes with high accuracy but also distinguishes 3 types of differential expression patterns which can reveal different functional mechanisms. We believe that DEsingle is a powerful tool that brings a new angle for the analysis of scRNA-seq data.

## References

Anders, S. and Huber, W. Differential expression analysis for sequence count data. Genome Biol 2010;11(10):R106.

Bacher, R. and Kendziorski, C. Design and computational analysis of single-cell RNA-sequencing experiments. Genome Biol 2016;17:63.

Boon, W.C., et al. Increasing cDNA yields from single-cell quantities of mRNA in standard laboratory reverse transcriptase reactions using acoustic microstreaming. Journal of Visualized Experiments Jove 2011 (53):e3144.

Brennecke, P., et al. Accounting for technical noise in single-cell RNA-seq experiments. Nat Methods 2013;10(11):1093–1095.

Delmans, M. and Hemberg, M. Discrete distributional differential expression (D3E)--a tool for gene expression analysis of single-cell RNA-seq data. BMC Bioinformatics 2016;17:110.

Garay, A.M., et al. On estimation and influence diagnostics for zero-inflated negative binomial regression models. Computational Statistics & Data Analysis 2011;55(3):1304–1318.

Islam, S., et al. Quantitative single-cell RNA-seq with unique molecular identifiers. Nat Methods 2014;11(2):163–166.

Kharchenko, P.V., Silberstein, L. and Scadden, D.T. Bayesian approach to single-cell differential expression analysis. Nat Methods 2014;11(7):740–742.

Macaulay, I.C. and Voet, T. Single cell genomics: advances and future perspectives. PLoS Genet 2014;10(1):e1004126.

Marinov, G.K., et al. From single-cell to cell-pool transcriptomes: stochasticity in gene expression and RNA splicing. Genome Res 2014;24(3):496–510.

Miao, Z. and Zhang, X. Differential expression analyses for single-cell RNA-Seq: old questions on new data. Quantitative Biology 2016;4(4):243–260.

Petropoulos, S., et al. Single-Cell RNA-Seq Reveals Lineage and X Chromosome Dynamics in Human Preimplantation Embryos. Cell 2016;165(4):1012–1026.

Wang, Y. and Navin, N.E. Advances and applications of single-cell sequencing technologies. Mol Cell 2015;58(4):598–609.

