## Supplementary Materials for "DEsingle for detecting three types of differential expression in single-cell RNA-seq data"

### 1 Supplementary Methods

#### 1.1 Using ZINB model to fit scRNA-seq data with excessive zeros

The zero percentages are very high for most genes in scRNA-seq data. Figure S1A shows the histogram of zero percentages of all expressed genes in a human embryonic scRNA-seq dataset (Petropoulos, et al., 2016). For RNA-seq read counts data, Negative Binomial (NB) distribution has been widely used in many methods (Robinson, et al., 2010). We adopted the zero-inflated Negative Binomial (ZINB) model to describe the read counts and the excessive zeros. We found the model fits scRNA-seq data well (Figure S1B).

The ZINB distribution is a mixture of constant zeros and a NB distribution with a mixing parameter $\theta$. The probability mass function (PMF) of ZINB distribution for read counts $N_g$ of gene $g$ in a group of cells is

$$\begin{aligned} P\left(N_g = n \mid \theta, r, p\right) &= \theta \cdot I\left(n=0\right) + \left(1-\theta\right) \cdot f_{NB}\left(r, p\right) \\ &= \theta \cdot I\left(n=0\right) + \left(1-\theta\right) \cdot \binom{n+r-1}{n} p^n \left(1-p\right)^r, \quad n = 0,1,2,... \end{aligned}$$

where $\theta$ is the proportion of constant zeros of gene $g$ in the group of cells, $I\left(n=0\right)$ is an indicator function which takes the value 1 for $n=0$ and 0 for $n \neq 0$, $f_{NB}$ is the PMF of the NB distribution, $r$ is the size parameter and $p$ is the probability parameter of the NB distribution. Note that the NB part can also have zero values. The observed zero values are the sum of constant zeros and zero values from the NB distribution.

#### 1.2 Modeling the mRNA capture procedure

Dropout zeros are mostly produced because of the low efficiency of mRNA capture in single cells (Boon, et al., 2011; Islam, et al., 2014; Macaulay and Voet, 2014; Marinov, et al., 2014). The mRNA products of some genes may be totally missed in the capturing procedure (Kharchenko, et al., 2014; Pierson and Yau, 2015; Wang and Navin, 2015). For convenience, we use "mRNA capture procedure" to refer to both the mRNA capturing in the reverse transcription step and the cDNA capturing in the cDNA amplification step. We modeled this procedure and studied its impact on the ZINB distribution.

Let $m_{ij}$ denote the original transcript copy number of gene $i$ in cell $j$, and let $n_{ij}$ denote the number of captured transcript copies of gene $i$ in cell $j$. The true $m_{ij}$ is unknown and the observed $n_{ij}$ is the result of sampling from existing transcripts with the mRNA capture procedure. We use a parameter $\beta \in [0\%, 100\%]$ to denote the efficiency of the mRNA capture procedure. Assume $m_{ij} \sim ZINB(\theta, r, p)$. Under the random capture assumption that all transcripts are captured with the same probability $\beta$, we can prove that $n_{ij} \sim ZINB(\theta, r, p^*)$ with $p^* = \frac{p\beta}{1-p+p\beta} \le p$ (Figure S1C), as described in following.

Proof:

Suppose $\{m_{ij}\}_i$ is from a unknown natural distribution $F$ over the cell population. Assume $F$ follows a ZINB distribution $F = \theta \cdot \delta_0 + (1-\theta) \cdot NB(r, p)$. Because the probability mass function (PMF) of NB is

$$f(x = k \mid r, p) = \binom{k+r-1}{k} p^k (1-p)^r,$$

for $k \in \{0, 1, 2, 3, ...\}$, $r \in (0, +\infty)$ and $p \in (0, 1)$, the PMF of $m_{ij}$ is

$$f(m_{ij} = k \mid \theta, r, p) = \begin{cases} \theta + (1-\theta)(1-p)^r, & if\ k = 0 \\ (1-\theta)\binom{k+r-1}{k} p^k (1-p)^r, & if\ k > 0 \end{cases}$$

And we have for all r and p that

$$\sum_{k=0}^{\infty} f(x = k \mid r, p) = \sum_{k=0}^{\infty} \binom{k+r-1}{k} p^k (1-p)^r = 1.$$

Under the random capture assumption that all transcripts are captured with the same probability $\beta$, we further have

$$n_{ij} \mid m_{ij} \sim Binomial(m_{ij}, \beta),$$

$$f\left(n_{ij}=k \mid m_{ij},\beta\right)=\binom{m_{ij}}{k}\cdot\beta^{k}\cdot(1-\beta)^{m_{ij}-k}.$$

Then, we have

$$\begin{aligned}f\left(m_{ij},n_{ij} \mid \theta,r,p,\beta\right)&=f\left(m_{ij} \mid \theta,r,p\right)\cdot f\left(n_{ij} \mid m_{ij},\beta\right)\\&=\theta\cdot I\left(m_{ij}=0\right)+(1-\theta)\binom{m_{ij}+r-1}{m_{ij}}p^{m_{ij}}(1-p)^{r}\binom{m_{ij}}{n_{ij}}\beta^{n_{ij}}(1-\beta)^{m_{ij}-n_{ij}}\end{aligned}$$

and

$$f\left(n_{ij} \mid \theta,r,p,\beta\right)=\sum_{k=0}^{\infty}f\left(n_{ij},m_{ij}=k \mid \theta,r,p,\beta\right).$$

1) When $n_{ij}=0$,

$$\begin{aligned}&f\left(n_{ij}=0 \mid \theta,r,p,\beta\right)\\&=\sum_{k=0}^{\infty}f\left(n_{ij}=0,m_{ij}=k \mid \theta,r,p,\beta\right)\\&=\theta+(1-\theta)(1-p)^{r}+\sum_{k=1}^{\infty}(1-\theta)\binom{k+r-1}{k}p^{k}(1-p)^{r}(1-\beta)^{k}\\&=\theta+(1-\theta)(1-p)^{r}+(1-\theta)(1-p)^{r}\frac{\sum_{k=0}^{\infty}\binom{k+r-1}{k}\left[p(1-\beta)\right]^{k}\left[1-p(1-\beta)\right]^{r}-\left[1-p(1-\beta)\right]^{r}}{\left[1-p(1-\beta)\right]^{r}}\\&=\theta+(1-\theta)\left[1-\frac{p\beta}{1-p+p\beta}\right]^{r}\end{aligned}$$

2) When $n_{ij}=t,\left(t=1,2,3,...\right)$,

$$\begin{aligned}&f\left(n_{ij}=t \mid \theta,r,p,\beta\right)\\&=\sum_{k\geq t}f\left(n_{ij}=t,m_{ij}=k \mid \theta,r,p,\beta\right)\\&=\sum_{k\geq t}(1-\theta)\binom{k+r-1}{k}p^{k}(1-p)^{r}\binom{k}{t}\beta^{t}(1-\beta)^{k-t}\\&=\frac{(t+r-1)!(1-\theta)(1-p)^{r}}{(r-1)!t!}\left(\frac{\beta}{1-\beta}\right)^{t}\left[p(1-\beta)\right]^{t}\sum_{k=0}^{\infty}\frac{(k+t+r-1)!}{(t+r-1)!k!}\left[p(1-\beta)\right]^{k}\\&=\binom{t+r-1}{t}(1-\theta)(1-p)^{r}(p\beta)^{t}\frac{\sum_{k=0}^{\infty}\frac{(k+t+r-1)!}{(t+r-1)!k!}\left[p(1-\beta)\right]^{k}\left[1-p(1-\beta)\right]^{t+r}}{\left[1-p(1-\beta)\right]^{t+r}}\\&=(1-\theta)\binom{t+r-1}{t}\left[\frac{p\beta}{1-p+p\beta}\right]^{t}\left[1-\frac{p\beta}{1-p+p\beta}\right]^{r}\end{aligned}$$

Therefore, we proved that the PMF of $n_{ij}$ is

$$f\left(n_{ij}=t\mid\theta,r,p^*\right)=\begin{cases}\theta+(1-\theta)\cdot\left(1-p^*\right)^r, & if\ t=0\\(1-\theta)\binom{t+r-1}{t}\left(p^*\right)^t\left(1-p^*\right)^r, & if\ t>0\end{cases}, \text{ where } p^*=\frac{p\beta}{1-p+p\beta}.$$

This proves that, if the true copy number $m_{ij}$ follows a ZINB distribution $m_{ij}\sim ZINB(\theta,r,p)$, after the mRNA capture step with efficiency $\beta$, the observed $n_{ij}$ is still a ZINB distribution. And the new distribution becomes $n_{ij}\sim ZINB(\theta,r,p^*)$, where $p^*=\frac{p\beta}{1-p+p\beta}$. The other parameters especially the $\theta$ is not changed.

Since $p-p^*=p-\frac{p\beta}{1-p+p\beta}=\frac{p-p^2+p^2\beta-p\beta}{1-p+p\beta}=\frac{p(1-p)(1-\beta)}{1-p+p\beta}\geq 0$, we have $p^*\leq p$.

The two major findings are:

1) Parameter $p$ in the ZINB model becomes $p^*=\frac{p\beta}{1-p+p\beta}\leq p$ after the capture procedure with efficiency $\beta$. The mean and the variance of the NB part of the ZINB model are $\frac{pr}{1-p}$ and $\frac{pr}{(1-p)^2}$, respectively. Therefore, when $p$ becomes smaller, both the mean and the variance of the NB part become smaller. This means that the NB distribution shifts toward zero after the capture procedure, and more zeros in the observed data will be from the NB part of the model. Figure S1D shows examples of theoretical ZINB distribution of original data ($\beta$ = 100%) and observed data with different capture efficiencies ($\beta$ = 30%, 20% and 10%).
2) The parameter $\theta$ is unchanged after the mRNA capture procedure, which means that the proportion of constant zeros of a gene among a group of cells is unchanged. For the original ideal data ($\beta$ = 100%), only cells with no transcript of the gene have zero values (assuming sufficiently deep sequencing). With imperfect capture efficiency, observed zeros become the mixture of real zeros and dropout zeros. Because $\theta$ is unchanged by the capture procedure, the $\theta$ estimated from observed data can be used to measure the proportion of real zeros in the original data. This is important for assessing the expression status of a gene among cells, although it cannot tell exactly which zero is real or dropout.

#### 1.3 Normalization

DEsingle integrates a modified median normalization method originated from DESeq to normalize the read counts data (Anders and Huber, 2010). Let $n_{ij}$ denote the read counts of gene $i$ in cell $j$, then the size factor of cell $j$ is estimated by $\hat{s}_j = \underset{i}{median} \frac{n_{ij}}{\left(\prod_{v=1}^{m} n_{iv}\right)^{1/m}}$, where $n_{iv} \neq 0$, namely only the non-zero read counts of each gene are used. Finally the normalized read counts of gene $i$ in cell $j$ is calculated by $n_{ij} / \hat{s}_j$.

#### 1.4 Detecting DE genes by comparing two ZINB populations

Detecting DE genes between two groups of cells using ZINB model is equivalent to testing the heterogeneity of two ZINB populations (Garay, et al., 2011; Rodrigues, 2006; Tse, et al., 2009). Each population is characterized by a ZINB model with three parameters. When any of the parameters of two models has significant difference, we consider the gene as differentially expressed. We use $\{n_{ij_1}\}$ and $\{n_{ij_2}\}$ to denote the read counts of gene $i$ in cell $j$ of group 1 and of group 2, respectively, and use the following 3 steps to test the difference of the two populations:

1) Calculate Maximum Likelihood Estimation (MLE) of the two ZINB populations' parameters $\hat{\boldsymbol{\theta}} = \left(\hat{\theta}_1, \hat{r}_1, \hat{p}_1, \hat{\theta}_2, \hat{r}_2, \hat{p}_2\right)$ with Expectation-Maximization (EM) algorithm (Dempster, et al., 1977) using $\{n_{ij_1}\}$ and $\{n_{ij_2}\}$ respectively.
2) Calculate constrained MLE of the two ZINB populations' parameters $\hat{\boldsymbol{\theta}}_0 = \left(\hat{\theta}_{1,0}, \hat{r}_{1,0}, \hat{p}_{1,0}, \hat{\theta}_{2,0}, \hat{r}_{2,0}, \hat{p}_{2,0}\right)$ under the null hypothesis $H_0 : \theta_1 = \theta_2, r_1 = r_2, p_1 = p_2$ using $\{n_{ij_1}\}$ and $\{n_{ij_2}\}$, respectively. It is equivalent to calculate the unconstrained MLE using $\{n_{ij_1}\}$ and $\{n_{ij_2}\}$ together.
3) Hypothesis testing of $H_0$. According to Wilks Theorem (Wilks, 1938), under the null hypothesis $H_0$, the $\chi^2_{LR1}$ statistics follows a $\chi^2_3$ distribution,

$$\chi^2_{LR1} = -2\log\lambda(n) = -2\log\frac{\sup_{\Theta_0} L(\boldsymbol{\theta} \mid \mathrm{n})}{\sup_{\Theta} L(\boldsymbol{\theta} \mid \mathrm{n})} = -2\left[l\left(\hat{\boldsymbol{\theta}}_0 \mid n\right) - l\left(\hat{\boldsymbol{\theta}} \mid n\right)\right] \sim \chi^2_3$$

where $\lambda(n)$ is the likelihood ratio statistic, $L(\boldsymbol{\theta} \mid \mathrm{n})$ is the likelihood function, $l(\boldsymbol{\theta} \mid n)$ is the log-likelihood function. Then the hypothesis testing of $H_0$ is conducted using the $\chi^2_{LR1}$ statistics.

#### 1.5 Subdividing three types of DE genes

DEsingle can subdivide three types of DE genes between single cell groups according to patterns of differences of parameters. The testing against the null hypothesis $H_0 : \theta_1 = \theta_2, r_1 = r_2, p_1 = p_2$ detects all genes that are differentially expressed between the two groups. We use another two hypothesis tests $H_{20}: \theta_1 = \theta_2$ and $H_{30}: r_1 = r_2, p_1 = p_2$ to characterize the patterns of the found DE genes.

The first type is DE genes with $H_0$ and $H_{20}$ rejected but with $H_{30}$ accepted (Table 1). This means that there is a significant difference in the number of cells with real zero values of the gene between the two groups of cells, but the expression in the remaining cells shows no significant difference. We call this type of genes as DEs genes or genes with different expression statuses. The second type is genes with $H_0$ and $H_{30}$ rejected but $H_{20}$ accepted (Table 1). They have differential expression abundances between the two groups, but do not show significant difference in the proportion of real zero values. We call this type as DEa genes. The third type is genes that have both different expression statuses and differential expression abundances. We call this type of genes as DEg genes or genes with general differential expression. Most of them are genes with $H_0$ rejected and also both $H_{20}$ and $H_{30}$ rejected. There may also be genes with $H_0$ rejected but both $H_{20}$ and $H_{30}$ accepted at the same significance level (Table 1).

We further breakdown the 3 types into 6 sub-categories according to the direction of the differences. For a DEs gene, there are significantly more cells with this gene "on" in the group with smaller $\theta$ (group 1) than in the other group (group 2). In other words, the gene tends to be turned on in group 1 and turned off in group 2. We name the gene as "DEs-on" in group 1 and "DEs-off" in group 2 for the convenience of future discussion. For a DEa gene, we call it as "DEa-up" in group 1 and "DEa-down" in group 2 if the mean of the NB part $\mu_{NB}$ is higher in group 1. For a DEg gene, we call it as "DEg-up" in the group with higher $\mu_{NB}(1-\theta)$, and call it as "DEg-down" in the other group. Table 1 provides a summary on the 3 types of DE genes and the 6 sub-categories.

**Table 1. Sub-categories of DE genes.** $\mu_{NB} = pr/(1-p)$, which is the mean of the ZINB model's NB part.

| DE type | DE parameters | Null Hypotheses Rejection | | | Conditions for "DEs-on/DEa-up/DEg-up" in group 1 |
|---|---|---|---|---|---|
| | | H0 | H20 | H30 | |
| Different expression status (DEs) | θ | Sig. | Sig. | Not sig. | $\theta_1 \le \theta_2$ |
| Differential expression abundance (DEa) | r or p | Sig. | Not sig. | Sig. | $\mu_{NB_1} \ge \mu_{NB_2}$ |
| General differential expression (DEg) | θ, and r or p | Sig. | Both sig. or both not sig. | | $\mu_{NB_1}(1-\theta_1) \ge \mu_{NB_2}(1-\theta_2)$ |

#### 1.6 Statistical test

Under the null hypothesis $H_0$, the $\chi^2_{LR1}$ statistics follows a $\chi^2_3$ distribution. So the p-value of $H_0$ could be derived by $p_{LR1} = 1 - F_{\chi^2_3}\left(\chi^2_{LR1}\right)$, where $F_{\chi^2_3}(x) = P\left(\chi^2_3 \le x\right)$. In other words, $F_{\chi^2_3}(x)$ is the cumulative distribution function of a chi-square distribution of freedom 3.

Similarly, we can derive the p-value of null hypothesis $H_{20}$ and $H_{30}$. For the multiple test of all the genes in one DE analysis, DEsingle gives the adjusted p-values to control the false discovery rate (Benjamini and Hochberg, 1995).

#### 1.7 Simulation study

We designed a series of simulation data to study the performance of the proposed method. We used the simulation method in (Delmans and Hemberg, 2016; Pierson and Yau, 2015) to generate scRNA-seq simulation data. In brief, the simulated data was generated by randomly sampling from a Poisson-Beta distribution (Delmans and Hemberg, 2016), which is originated from the transcriptional bursting model (Gillespie, 1976). The data were produced with the following procedures: Firstly, draw a variable $c$ from a Beta distribution with parameters $\alpha$ and $\beta$, namely $c \sim Beta(\alpha, \beta)$. Secondly, randomly sample a number from the Poisson distribution with parameter $\lambda = c\gamma$. Parameters $\alpha$, $\beta$ and $\gamma$ were picked from the parameters list of (Delmans and Hemberg, 2016) which was generated from a real scRNA-seq dataset (Islam, et al., 2011).

We applied the dropout model introduced by Pierson and Yau to add dropout events to the simulated data (Pierson and Yau, 2015). Specifically, let $x_{ij}$ denote the expression level of gene $i$ in cell $j$, and let $\mu$ denote the mean of non-zero expression level (log read counts) of gene $i$ across cells. The dropout rate of gene $i$ is modeled as $p_0 = \exp\left(-\lambda\mu^2\right)$, where $\lambda$ is the exponential decay parameter and is shared across genes. We use $\lambda = 0.1$ in our simulation as recommended. Then the expression level $y_{ij}$ of gene $i$ in cell $j$ after adding dropout events is denoted as $y_{ij} = \begin{cases} x_{ij}, & \text{if } h_{ij} = 0, \\ 0, & \text{if } h_{ij} = 1, \end{cases}$, where $h_{ij} \mid x_{ij} \sim \text{Bernoulli}\left(p_0\right)$.

Using this strategy, we simulated two types of scRNA-seq data: those with and without additional dropout events. For each type of data, we simulated datasets with $10 \times 2$, $50 \times 2$, $100 \times 2$ and $200 \times 2$ cells, each with 10,000 simulated genes. We got 8 sets of simulation data in this way, 4 with additional dropout events and 4 without.

For each dataset, we set half of the genes as DE genes that have fold changes greater than 1.5 in any of the three parameters between the two groups.

We applied the proposed DEsingle and 7 other methods on the 8 sets of simulation data. The compared methods are BPSC (Vu, et al., 2016), D3E (Delmans and Hemberg, 2016), monocle (Trapnell, et al., 2014), SCDE (Kharchenko, et al., 2014), DESeq2 (Love, et al., 2014), edgeR (Robinson, et al., 2010) and DEGseq (Wang, et al., 2010). Four of the methods, namely BPSC, D3E, monocle and SCDE, are specially designed for scRNA-seq data. The other three methods, DESeq2, edgeR and DEGseq, are traditional DE analysis methods developed for bulk RNA-seq data, but have also been widely applied on single-cell data. The parameters settings of each method in the experiments were provided in section 4.7.

#### 1.8 Case study on real data

We applied DEsingle on a public scRNA-seq dataset of human preimplantation embryonic cells (Petropoulos, et al., 2016). We conducted a systematic comparison on the gene expression of the 81 cells from E3 and that of the 190 cells from E4 in this dataset. We used the mapped raw read counts table provided by the authors as input to DEsingle to detect and analyze the three types of DE genes between E3 and E4 cells.

#### 1.9 Parameters settings of DE analysis methods

BPSC(v.0.99.0): The analysis is conducted as the examples shown in BPSC package introduction.

D3E (Latest commit 6727adf on 21 Oct 2015): All the parameters are as suggested in the example of D3E's wiki pages, except '-m 0', because mode 1 runs too slow for the large scale experiments.

monocle (v.1.99.0): The parameter expressionFamily for function newCellDataSet is 'expressionFamily=negbinomial()'. The parameter cores for function differentialGeneTest is 'cores = 8'. Other parameters are as vignette of monocle suggested.

SCDE (v.1.99.1): The parameters of function clean.counts are 'min.reads = 0' and 'min.detected = 0'. We set paramrter 'n.cores = 8' for function scde.error.models and function scde.expression.difference. Other parameters are as suggested in SCDE's tutorials.

DESeq2 (v.1.18.1): All the parameters setting are as suggested in the Quick start part of DESeq2's documentation.

edgeR (v.3.8.6): All the parameters setting are as suggested by the documentation of edgeR.

DEGseq (v.1.20.0): The parameter used in DEGexp is ‘method="MARS"’ and other parameters are as the examples of DEGexp shown in documentation of DEGseq.

### 2 Supplementary Results

#### 2.1 Three types of DE genes between E3 and E4 of human embryonic cells

We applied DEsingle to compare the 81 embryonic day 3 (E3) cells to the 190 embryonic day 4 (E4) cells from the human preimplantation embryonic dataset (see Methods) (Petropoulos, et al., 2016). These are two days that the embryonic cells undergo dramatic changes. DEsingle reported a total of 7,560 DE genes at the significance level of p-value<0.05 after Bonferroni correction. The majority of them, 5,685 genes (75.2%), belong to the DEg type. Among them, 2,909 genes are DEg-up and 2,776 genes are DEg-down in E3 cells compared to E4 cells. There are another 1,147 genes (15.2%) that are of the DEa type, with 670 DEa-up genes and 477 DEa-down genes in E3 compared to E4. Besides these genes of the more conventional sense of differential expression, DEsingle reported 728 significant DEs genes (9.6%), with 333 genes of DEs-on in E3 and 395 genes of DEs-on in E4. We also applied other methods including BPSC, D3E, monocle, SCDE, DESeq2, edgeR and DEGseq on this data as a comparison. We found that most methods including BPSC, monocle, SCDE, DESeq2 and edgeR can detect only a small proportion of the DEs genes when the reported DE gene numbers are at the same level with that reported by DEsingle. Methods like D3E and DEGseq can recover almost all the DEs genes at the cost of reporting huge numbers of DE genes (17,889 by D3E and 17,801 by DEGseq).

Figure S3A shows the heatmap of top 500 genes of each of the 3 types of DE genes (details of the genes are shown in Additional file 1: Table S1). We can see that DE genes of the same type tends to be clustered together, and each type has its distinct expression pattern in E3 and E4. Figure S3B shows expression histograms of 3 example genes in E3 cells and E4 cells, illustrating the patterns of differences in DEs, DEa and DEg genes.

To explore the potential functional roles of each type of DE genes, we performed Gene Ontology (GO) enrichment analysis on the top 500 DE genes of each type using DAVID (Huang da, et al., 2009). The 1,500 DE genes were divided into 6 sub-categories according to Table 1. They include 238 DEs-on, 274 DEa-up and 232 DEg-up genes in E3, and 262 DEs-on, 226 DEa-up and 268 DEg-up genes in E4. Figure S4A shows the overlap between the enriched GO terms from the 6 sub-categories of DE genes. Table S2 (Additional file 2) lists all enriched GO terms with some key functions highlighted in Figure S4B.

From Figure S4A, we can see that there are more enriched GO terms in DEa and DEg genes than in DEs genes. We can infer that most of the transcriptome differences between E3 and E4 can be captured by the conventional type of differential expression. However, DEs genes compose a noticeable number of the DE genes, which implies that there are genes that are switched on or off between E3 and E4. The GO terms enriched by these DEs genes have no intersection with the GO terms enriched by the other DE genes (Figure S4A). Specifically, there are 9 GO functions that are uniquely enriched by the DEs-on genes in E3, and 2 GO functions uniquely enriched by the DEs-on genes in E4. The regulation and function of DEs genes may involve different pathways with the DEa or DEg genes.

DEs-on genes of E3 are enriched for Biological Process (BP) GO terms of cell adhesion, extracellular matrix organization, Cellular Component (CC) GO terms of integral component of plasma membrane, plasma membrane, basement membrane, cell junction, and Molecular Function (MF) GO terms of calcium ion binding, extracellular ligand-gated ion channel activity (Figure S4B; Additional file 2: Table S2). It is known that E3 is around the 8-cell stage of preimplantation embryos (Niakan, et al., 2012), during which cell compaction occurs and functional gap junctions are formed (Brison, et al., 2014; Fleming, et al., 2001). Cell compaction is initiated by the E-cadherin mediated cell adhesion (Adams, et al., 1998; Fleming, et al., 2001; Larue, et al., 1994; Li, et al., 2009). Genes related to cytoskeletal, cell junction and cell adhesion are also involved in this process (Cui, et al., 2007). In the DEsingle results, we see that the enriched GO terms of E3 DEs-on genes are all associated with the cell compaction process. For example, calcium ion binding and extracellular ligand-gated ion channel activity are known to play an important role to the function and interactions of E-cadherin (Kim, et al., 2011); extracellular matrix organization, integral component of plasma membrane, plasma membrane, basement membrane and cell junction are essential to the formation of gap junctions and cell adhesion (Brison, et al., 2014; Cui, et al., 2007; Fleming, et al., 2001; Li, et al., 2009). Take the E3 DEs-on genes ICAM1 (intercellular adhesion molecule 1) and ICAM5 (intercellular adhesion molecule 5, shown in Figure S3B) as examples. They show up in 4 of the above GO terms. The zero expression ratios of gene ICAM1 and ICAM5 in E3 cells are 28.4% and 45.3%, respectively, while the ratios in E4 cells are 73.6% and 98.2%, respectively. In other words, these two genes are actively expressed in most of E3 cells but are turned off in almost all E4 cells. We can infer that genes related to cell compaction are active in E3 but are turned off gradually in E4 after compaction is finished and morula is formed (Figure S4B). Recent studies have shown that, although the initiation of cell compaction occurred from 4-cell to 16-cell stage (about late embryonic day 2 to embryonic day 4), most human embryos (86.1%) initiated compaction at the 8-cell stage or later (around E3 to E4 stage), with initiation at the

8-cell stage (E3) being most frequent (Iwata, et al., 2014). This confirms the inference on the DEs genes and also explains why some compaction related genes are still expressed in very few E4 cells.

The E4 DEs-on genes are enriched for the BP GO term of positive regulation of collagen biosynthetic process (Figure S4B; Additional file 2: Table S2). According to a recent study of Petropoulos et al. (Petropoulos, et al., 2016), human preimplantation embryonic cells will differentiate into three lineages of epiblast (EPI), primitive endoderm (PE) and trophectoderm (TE) simultaneously at E5. Rasmussen et al. had shown that collagen could improve the differentiation of human embryonic cells towards endoderm (Rasmussen, et al., 2015), which implies the synthesis of collagen in some of the morula cells at E4 is very likely associated with the differentiation of PE cells at E5. This explains why the positive regulation of collagen biosynthetic process is enriched in E4 DEs-on genes, and also verifies the different expression statuses of the related genes between E3 and E4 cells. Besides, genes TGFB1 and TGFB3 are also found in the E4 DEs-on category. The zero expression ratios of gene TGFB1 and TGFB3 in E3 cells are 87.9% and 92.5%, respectively, but are reduced in E4 cells to 20.6% and 33.8%, respectively. In other words, they are turned on in a large proportion of cells in E4 compared to E3. The TGF-β ligands encoded by TGFB1 and TGFB3 are triggers of the TGF-β signaling pathway, which are enriched in EPI cells in E5 (Petropoulos, et al., 2016).

For the other types of DE genes, E3 DEa-up genes are enriched for BP GO terms of mRNA splicing via spliceosome, mRNA 3'-end processing, RNA splicing, mRNA export from nucleus, RNA export from nucleus, mRNA processing, and CC GO terms of nucleoplasm, nucleus, nucleolus (Figure S4B; Additional file 2: Table S2). E3 DEg-up genes also are enriched in CC GO terms of nucleus and nucleoplasm. These GO terms are consistent with the process of embryonic genome activation (EGA) that occurs at E3 (Niakan, et al., 2012; Petropoulos, et al., 2016), which brings the burst of transcription (Christians, et al., 1995) in nucleus (Nelson and Cox, 2005; Weaver, 2008) simultaneously (Figure S4B).

E4 DEa-up genes are enriched in BP GO terms of translational initiation, translation, cytoplasmic translation, CC GO terms of ribosome, cytosol, cytoplasm, and MF GO terms of translation initiation factor activity (Figure S4B; Additional file 2: Table S2). These enriched GO terms are in agreement with the fact that translation occurs on the ribosomes in the cytoplasm (Nelson and Cox, 2005; Weaver, 2008), and the translation peak of EGA is delayed (Hamatani, et al., 2004; Nothias, et al., 1995) to E4, which is called uncoupling of transcription and translation during EGA (Schultz, 2002).

Except for the GO terms related to translation, E4 DEg-up genes are also enriched for BP GO terms of mitochondrial translational elongation, mitochondrial respiratory chain complex I assembly, mitochondrial electron transport NADH to ubiquinone, mitochondrial translation, mitochondrial ATP synthesis coupled proton transport, protein targeting to mitochondrion, and CC GO terms of mitochondrial inner membrane, mitochondrion, mitochondrial respiratory chain complex I, and mitochondrial ribosome (Figure S4B; Additional file 2: Table S2). All these enriched GO terms are directly or indirectly related to mitochondrion. Dumollard et al. have reported that before the compaction is complete, pyruvate is metabolized by mitochondria whereas glucose is not, and that after compaction, glucose gradually becomes the major substrate for energy supply instead of pyruvate in embryonic cells (Dumollard, et al., 2007) (Figure S4B). Moreover, Wilding et al. have showed that the aerobic respiration is upregulated and the percentage of glucose metabolised through aerobic respiration rises dramatically during this period (Wilding, et al., 2009). These reports as well as our observations imply that the major metabolic mode of mitochondrion is changed as early as from E3 to E4 and the mitochondrion becomes more active at E4. It is reasonable to infer that the changes of the metabolic mode and activity of mitochondrion are related to the translation peak in E4 which requires more energy than E3.

### 3 Supplementary Discussion

Differential expression analysis is an important aspect of scRNA-seq data analyses (Kharchenko, et al., 2014; Stegle, et al., 2015; Wang and Navin, 2015). The excessive amount of zero values composed of the mixture of real zeros and dropout zeros pose a great challenge to DE genes detection (Poirion, et al., 2016; Saliba, et al., 2014; Stegle, et al., 2015), but also provide opportunities for detecting different patterns of differential expression at single-cell level. Existing DE analysis methods are not capable of distinguishing the two types of zeros (Delmans and Hemberg, 2016; Kharchenko, et al., 2014; Trapnell, et al., 2014; Vu, et al., 2016), and may therefore miss important indicators of gene expression status in the cells.

We developed a method called DEsingle that can estimate the proportion of the two kinds of zeros and then detect the DE genes by taking account differences in this proportion. We studied the mRNA capture procedure using the ZINB model and proved the effects of the mRNA capture procedure on the model parameters. These results provide a way to estimate the proportion of real zero expressions of a gene in a group of cells. With this model, we can not only detect differentially expressed genes at higher accuracy, but also subdivide differential expression into three types according to the pattern of the differences. The types of DEg and DEa are close to the conventional understanding of differential expression, in the sense that a DE gene tend to be expressed significantly higher in one group of samples than in the other group of samples. The type DEs or different expression status addresses a unique phenomenon in single-cell transcriptomes. The transcription of a gene in a single cell is a discrete on-off process. The DEs genes identified by DEsingle have a significantly higher proportion of being on (or off) in one group of cells than in the other group of cells. This question has not been considered in existing literature on differential expression analysis.

We used simulation experiments to compare the performance of DEsingle with representative existing methods for DE analysis on single-cell data. Results showed the advantage of DEsingle in detecting DE genes with high accuracy. We applied the method on a human preimplantation embryonic cell scRNA-seq dataset to study the biological significance of the proposed method, especially the subdivision of the three types of differential expression. Results showed that DEsingle can reveal significant genes and functional pathways that have significant changes in their expression between two days of embryonic cell development. Especially, the identified DEs genes revealed functions that tend to be switched on or off in the majority of cells from day 3 to day 4 along the embryonic development. The subdivision of DE genes into subtypes provides information at finer resolution for understanding changes of transcription activities and their regulations of a biological process.

### 4 Supplementary Figures

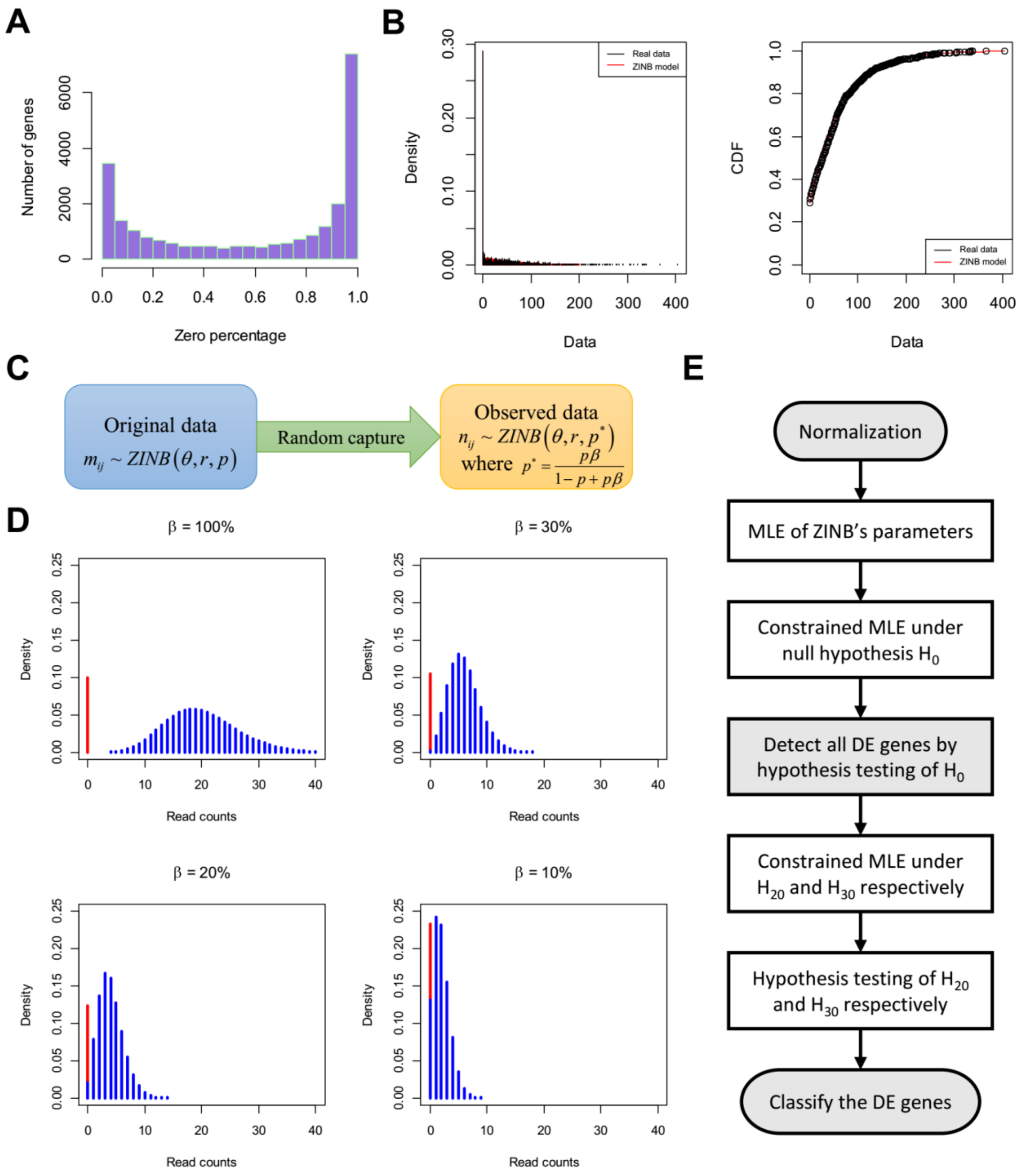

**Figure S1. ZINB model for scRNA-seq data and workflow of DEsingle.**

(A) Histogram of zero percentages of all expressed genes in a real scRNA-seq dataset. (B) An example of ZINB model fitting for scRNA-seq data. The left panel shows the density fitting. The right panel shows the cumulative distribution function fitting. (C) The mRNA capture procedure transforms the original ZINB model to another ZINB model with only one parameter changed. (D) Theoretical ZINB distribution of a gene with different random capture efficiency $\beta$. The original data corresponds to the case of $\beta$ = 100%. The cases of $\beta$ =30%, 20% and 10% represent the observed data obtained after mRNA capture procedure with different efficiency.

The parameters of original ZINB distribution are $\theta$=0.1, $r$=20 and $p$=0.5. The red bars represent the probability density of real zero expression, which is comes from the $\theta$ parameter of the ZINB model; the blue bars represent the probability density of the NB part of the ZINB model. When $\beta$ decreases, the zero density from NB part (the blue bar at zero) becomes larger. (E) Workflow of DEsingle to detect and classify DE genes. Hypothesis testing of $H_0 : \theta_1 = \theta_2, r_1 = r_2, p_1 = p_2$ is used to detect all the DE genes; hypothesis testing of $H_{20} : \theta_1 = \theta_2$ and $H_{30} : r_1 = r_2, p_1 = p_2$ are used to classify the found DE genes.

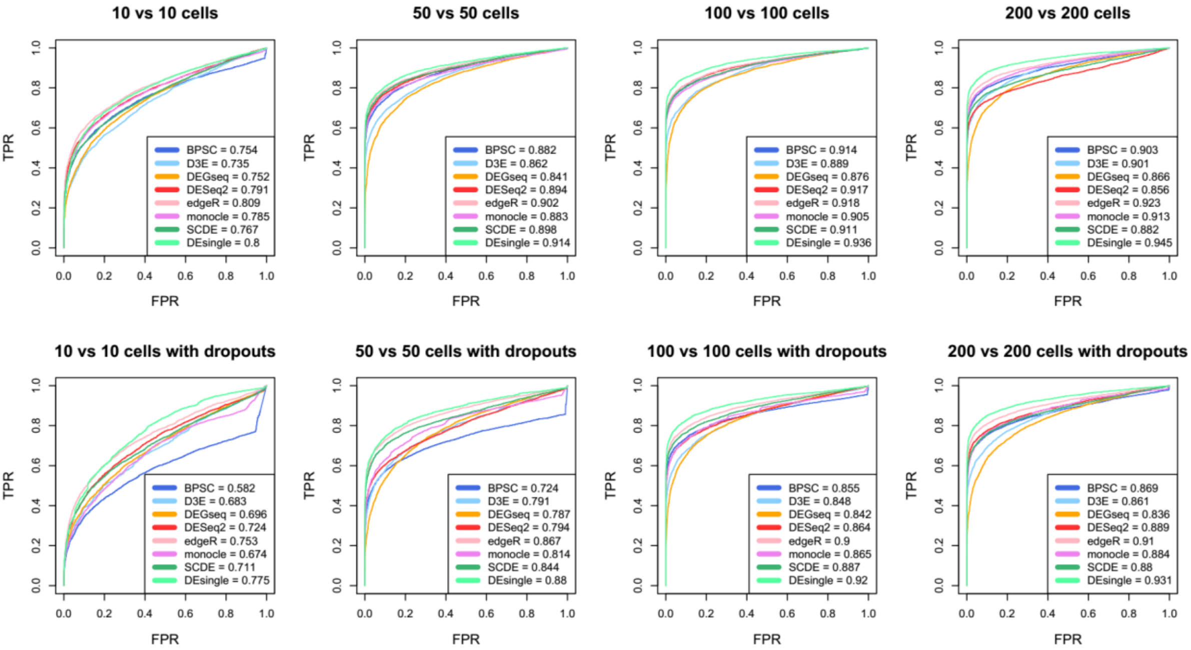

**Figure S2. Performances of DE analysis methods on simulation data.**

ROC curves and AUCs of 8 DE analysis methods on the 8 sets of simulated scRNA-seq data. The top 4 graphs are on simulation data without additional dropout events. The bottom 4 are on data with additional dropout events. The sample size for comparison is annotated above each graph. The AUC of each method is listed on bottom right corner of each graph. DEsingle has the best AUC in almost all plots except in the upper-left one where its AUC is slightly smaller than that of edgeR.

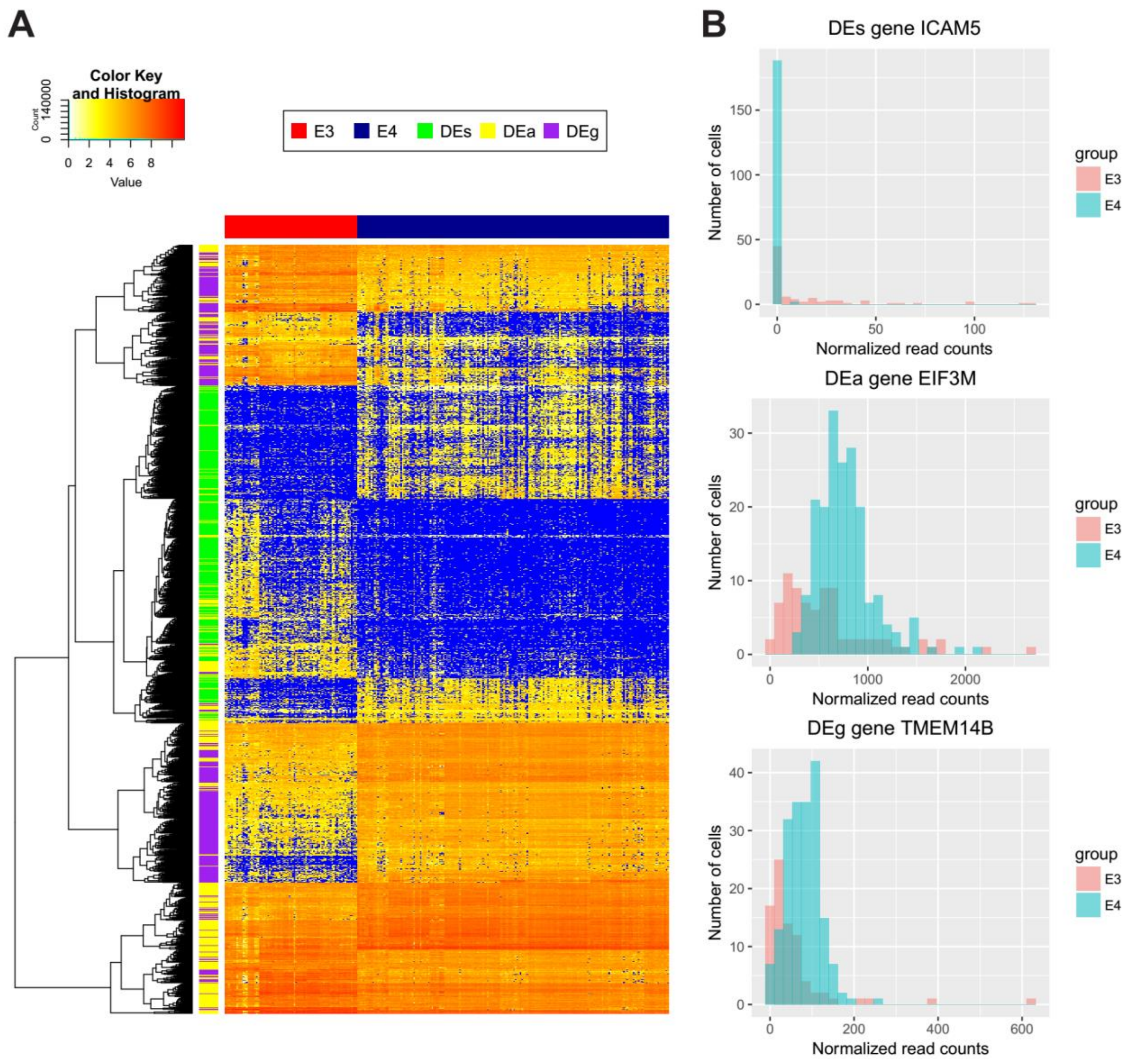

**Figure S3. Heatmap and example histograms of DE genes found by DEsingle.**

(A) Heatmap of expression levels (log(normalized counts + 1)) of top 500 genes of each type of DE genes found by DEsingle in the comparison of E3 versus E4 human preimplantation embryonic cells. Each row illustrates a gene and each column illustrates cell. The color blue is used to represent zero values and the non-zero values are coded by the gradual change of colors from light yellow to deep red. The genes are hierarchically clustered using complete-linkage clustering with Euclidean distance. (B) Histogram of expression levels of three example DE genes, belong to the types of DEs, DEa and DEg, respectively.

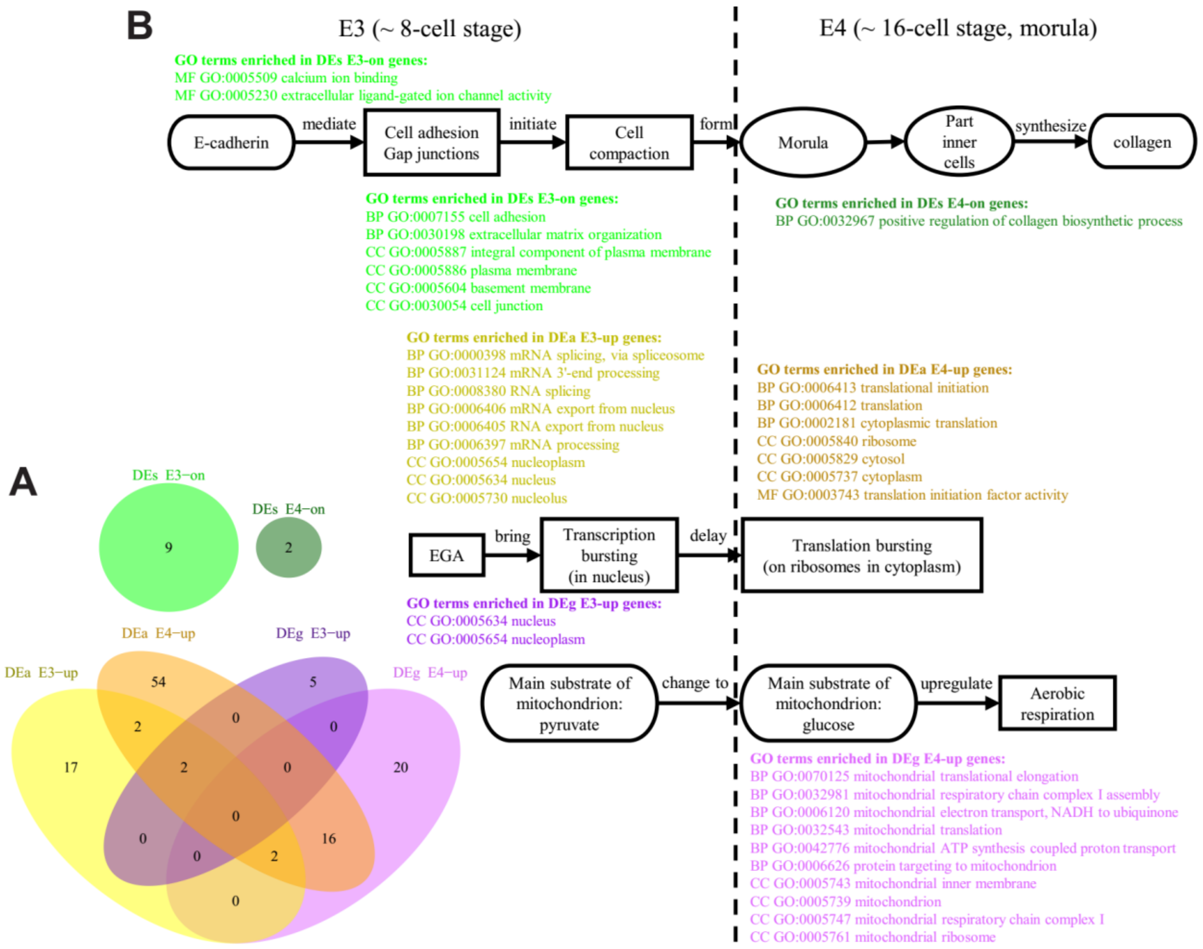

**Figure S4. Enriched GO terms and related biological processes of DE genes.**

(A) Venn diagram of the enriched GO terms in the 6 sub-categories of DE genes between embryonic day 3 (E3) to embryonic day 4 (E4) of the human preimplantation embryos. The GO terms enriched in DEs genes are unique and do not overlap with those enriched by other types of DE genes. (B) An illustrative diagram of the major biological processes from E3 to E4 and some significantly enriched GO terms along processes. EGA: embryonic genome activation.

### Additional files

Additional file 1: Table S1. Differential expression analysis of E3 and E4 cells with DEsingle.

Additional file 2: Table S2. GO enrichment analysis of DE genes using DAVID.

### List of abbreviations

scRNA-seq: Single-cell RNA sequencing; DE: Differential expression; ZINB: Zero-Inflated Negative Binomial; DEs: different expression status; DEa: different expression abundance; DEg: general differential expression; NB: Negative Binomial; PMF: Probability Mass Function MLE: Maximum Likelihood Estimation; ROC: Receiver Operating Characteristic; AUC: Area Under Curve; E3: embryonic day 3; E4: embryonic day 4; GO: Gene Ontology; BP: Biological Process; CC: Cellular Component; MF: Molecular Function; EPI: epiblast; PE: primitive endoderm; TE: trophectoderm; EGA: embryonic genome activation; EM: Expectation-Maximization.

### Availability of data and materials

The implemented R package DEsingle and its source code is available at https://github.com/miaozhun/DEsingle. The datasets generated and/or analyzed during the current study are available in the ArrayExpress repository, E-MTAB-3929 in https://www.ebi.ac.uk/arrayexpress/experiments/E-MTAB-3929/.

### References

Adams, C.L., *et al.* Mechanisms of epithelial cell–cell adhesion and cell compaction revealed by high-resolution tracking of E-cadherin–green fluorescent protein. *The Journal of cell biology* 1998;142(4):1105-1119.

Anders, S. and Huber, W. Differential expression analysis for sequence count data. *Genome Biol* 2010;11(10):R106.

Benjamini, Y. and Hochberg, Y. Controlling the false discovery rate: a practical and powerful approach to multiple testing. *Journal of the royal statistical society. Series B (Methodological)* 1995:289-300.

Boon, W.C., *et al.* Increasing cDNA yields from single-cell quantities of mRNA in standard laboratory reverse transcriptase reactions using acoustic microstreaming. *Journal of Visualized Experiments Jove* 2011(53):e3144.

Brison, D.R., Sturmey, R.G. and Leese, H.J. Metabolic heterogeneity during preimplantation development: the missing link? *Hum Reprod Update* 2014;20(5):632-640.

Christians, E., *et al.* Expression of the HSP 70.1 gene, a landmark of early zygotic activity in the mouse embryo, is restricted to the first burst of transcription. *Development* 1995;121(1):113-122.

Cui, X.S., *et al.* Transcription profile in mouse four-cell, morula, and blastocyst: Genes implicated in compaction and blastocoel formation. *Molecular reproduction and development* 2007;74(2):133-143.

Delmans, M. and Hemberg, M. Discrete distributional differential expression (D3E)--a tool for gene expression analysis of single-cell RNA-seq data. *BMC Bioinformatics* 2016;17:110.

Dempster, A.P., Laird, N.M. and Rubin, D.B. Maximum likelihood from incomplete data via the EM algorithm. *Journal of the royal statistical society. Series B (methodological)* 1977:1-38.

Dumollard, R., Duchen, M. and Carroll, J. The role of mitochondrial function in the oocyte and embryo. *Current topics in developmental biology* 2007;77:21-49.

Fleming, T.P., Sheth, B. and Fesenko, I. Cell adhesion in the preimplantation mammalian embryo and its role in trophectoderm differentiation and blastocyst morphogenesis. *Front Biosci* 2001;6(1):D1000-D1007.

Garay, A.M., *et al.* On estimation and influence diagnostics for zero-inflated negative binomial regression models. *Computational Statistics & Data Analysis* 2011;55(3):1304-1318.

Gillespie, D.T. A general method for numerically simulating the stochastic time evolution of coupled chemical reactions. *Journal of computational physics* 1976;22(4):403-434.
Hamatani, T., *et al.* Dynamics of global gene expression changes during mouse preimplantation development. *Developmental cell* 2004;6(1):117-131.
Huang da, W., Sherman, B.T. and Lempicki, R.A. Systematic and integrative analysis of large gene lists using DAVID bioinformatics resources. *Nat Protoc* 2009;4(1):44-57.
Islam, S., *et al.* Characterization of the single-cell transcriptional landscape by highly multiplex RNA-seq. *Genome Res* 2011;21(7):1160-1167.
Islam, S., *et al.* Quantitative single-cell RNA-seq with unique molecular identifiers. *Nat Methods* 2014;11(2):163-166.
Iwata, K., *et al.* Analysis of compaction initiation in human embryos by using time-lapse cinematography. *Journal of assisted reproduction and genetics* 2014;31(4):421-426.
Kharchenko, P.V., Silberstein, L. and Scadden, D.T. Bayesian approach to single-cell differential expression analysis. *Nat Methods* 2014;11(7):740-742.
Kim, S.A., *et al.* Calcium-dependent dynamics of cadherin interactions at cell–cell junctions. *Proceedings of the National Academy of Sciences* 2011;108(24):9857-9862.
Larue, L., *et al.* E-cadherin null mutant embryos fail to form a trophectoderm epithelium. *Proceedings of the National Academy of Sciences* 1994;91(17):8263-8267.
Li, C.-B., *et al.* Regulation of compaction initiation in mouse embryo. *Hereditas (Beijing)* 2009;31(12):1177-1184.
Love, M.I., Huber, W. and Anders, S. Moderated estimation of fold change and dispersion for RNA-seq data with DESeq2. *Genome biology* 2014;15(12):550.
Macaulay, I.C. and Voet, T. Single cell genomics: advances and future perspectives. *PLoS Genet* 2014;10(1):e1004126.
Marinov, G.K., *et al.* From single-cell to cell-pool transcriptomes: stochasticity in gene expression and RNA splicing. *Genome Res* 2014;24(3):496-510.
Nelson, D.L. and Cox, M.M. Principles of biochemistry. In.: New York: WH Freeman and Company; ISBN 0 7167 4339 6; 2005.
Niakan, K.K., *et al.* Human pre-implantation embryo development. *Development* 2012;139(5):829-841.
Nothias, J.-Y., *et al.* Regulation of gene expression at the beginning of mammalian development. *Journal of Biological Chemistry* 1995;270(38):22077-22080.
Petropoulos, S., *et al.* Single-Cell RNA-Seq Reveals Lineage and X Chromosome Dynamics in Human Preimplantation Embryos. *Cell* 2016;165(4):1012-1026.
Petropoulos, S., *et al.* Single-cell RNA sequencing: revealing human pre-implantation development, pluripotency and germline development. *Journal of internal medicine* 2016;280(3):252-264.
Pierson, E. and Yau, C. ZIFA: Dimensionality reduction for zero-inflated single-cell gene expression analysis. *Genome Biol* 2015;16:241.
Poirion, O.B., *et al.* Single-Cell Transcriptomics Bioinformatics and Computational Challenges. *Front Genet* 2016;7:163.
Rasmussen, C.H., *et al.* Collagen Type I Improves the Differentiation of Human Embryonic Stem Cells towards Definitive Endoderm. *PLoS One* 2015;10(12):e0145389.
Robinson, M.D., McCarthy, D.J. and Smyth, G.K. edgeR: a Bioconductor package for differential expression analysis of digital gene expression data. *Bioinformatics* 2010;26(1):139-140.
Rodrigues, J. Full Bayesian Significance Test for Zero-Inflated Distributions. *Communications in Statistics: Theory and Methods* 2006;35(2):299-307.
Saliba, A.E., *et al.* Single-cell RNA-seq: advances and future challenges. *Nucleic Acids Res* 2014;42(14):8845-8860.
Schultz, R.M. The molecular foundations of the maternal to zygotic transition in the preimplantation embryo. *Human Reproduction Update* 2002;8(4):323-331.
Stegle, O., Teichmann, S.A. and Marioni, J.C. Computational and analytical challenges in single-cell transcriptomics. *Nat Rev Genet* 2015;16(3):133-145.
Trapnell, C., *et al.* The dynamics and regulators of cell fate decisions are revealed by pseudotemporal ordering of single cells. *Nat Biotechnol* 2014;32(4):381-386.

Tse, S.K., *et al.* Testing homogeneity of two zero-inflated Poisson populations. *Biom J* 2009;51(1):159-170.
Vu, T.N., *et al.* Beta-Poisson model for single-cell RNA-seq data analyses. *Bioinformatics* 2016;32(14):2128.
Wang, L., *et al.* DEGseq: an R package for identifying differentially expressed genes from RNA-seq data. *Bioinformatics* 2010;26(1):136-138.
Wang, Y. and Navin, N.E. Advances and applications of single-cell sequencing technologies. *Mol Cell* 2015;58(4):598-609.
Weaver, R.F. Molecular Biology. McGraw-Hill; 2008.
Wilding, M., *et al.* Mitochondria and human preimplantation embryo development. *Reproduction* 2009;137(4):619-624.
Wilks, S.S. The large-sample distribution of the likelihood ratio for testing composite hypotheses. *The Annals of Mathematical Statistics* 1938;9(1):60-62.
